# A hyperthermophilic phage decoration protein suggests common evolutionary origin with Herpesvirus Triplex proteins and an anti-CRISPR protein

**DOI:** 10.1101/216911

**Authors:** Nicholas P. Stone, Brendan J. Hilbert, Daniel Hidalgo, Kevin T. Halloran, Brian A. Kelch

**Author notes:** Currently at Sanofi-Genzyme, Framingham, MA 01701.

## Abstract

Virus capsid proteins reproducibly self-assemble into regularly-shaped, stable shells that protect the viral genome from external environmental assaults, while maintaining the high internal pressure of the tightly packaged viral genome. To elucidate how capsids maintain stability under harsh conditions, we investigated the capsid components of a hyperthermophilic virus, phage P74-26. We determined the structure of a capsid protein gp87 and show that it has the same fold as trimeric decoration proteins that enhance the structural stability of capsids in many other phage, despite lacking significant sequence homology. We also find that gp87 is significantly more stable than its mesophilic homologs, reflecting the high temperature environment in which phage P74-26 thrives. Our analysis of the gp87 structure reveals that the core domain of the decoration protein is conserved in trimeric capsid components across numerous dsDNA viruses, including human pathogens such as Herpesviruses. Moreover, this core β-barrel domain is found in the anti-CRISPR protein AcrIIC1, which suggests a mechanism for the evolution of this broad spectrum Cas9 inhibitor. Our work illustrates the principles for increased stability of a thermophilic decoration protein, and extends the evolutionary reach of the core trimeric decoration protein fold.

## Introduction

Tailed bacteriophages (also known as Caudoviruses) are the most common biological entities on Earth [1,2]. Caudoviruses assemble viral particles by actively packaging the viral genome inside a preformed, self-assembled capsid structure [3]. Because the DNA fills the capsid to nearly crystalline density, the pressure inside the capsid head is estimated to be extremely high (>6 MPa) [4]. Thus, the capsid structure must withstand high internal strain, in addition to the challenges imposed by the fluctuating external environment.

The capsids of Caudoviruses are primarily comprised of a single major capsid protein (MCP) of the HK97 fold, which assembles into an icosahedral shell [5]. The MCP subunits loop through each other to build an interdigitated structure with high structural integrity. Proper construction of the capsid is directed by the portal ring that nucleates self-assembly of MCP, as well as a scaffolding protein that chaperones the MCP subunits to ensure the proper size and shape of the shell [6-9]. The MCP protein forms units called capsomers of five or six subunits, termed pentons or hexons, respectively. The capsid typically assembles with 11 pentons and a variable number of hexons, depending on the triangulation number of the virus shell. The capsid structure is altered during genome packaging, causing expansion of the shell, loss of the scaffolding protein, and a morphological change from nearly spherical to an icosahedral structure. The icosahedral shell is stable against the high internal pressure from the condensed DNA filling the capsid interior.

The three-fold vertices of the capsid icosahedron appear to be a weak point in the structure, and these viruses have evolved mechanisms to ensure stabilization of these vertices. For example, the siphovirus HK97 uses covalent linkages at the three-fold vertices that form a ‘chain-mail’ to stabilize the capsid structure [10]. On the other hand, many viruses instead use auxiliary capsid proteins to add structural integrity, in particular the trimeric decoration proteins that bind at the three-fold vertices of the capsid icosahedron [11]. Decoration proteins (also known as cementing proteins) play an important role in capsid stability and assembly. Cryo-electron microscopy (cryoEM) structures reveal that decoration proteins specifically interact with the three-fold axis by inserting its N-terminal region into a groove formed by neighboring capsomers [12]. Decoration proteins only bind to the expanded, icosahedral capsid because the sites for binding are occluded in the smaller, spherical procapsids [13]. Studies of the λ phage decoration protein revealed that binding of decoration protein trimers strengthens the capsid assembly against both increased temperature and mechanical deformation [11,14,15]. However, it is unclear how evolution modulates decoration protein-mediated stabilization. Moreover, it is unclear how decoration proteins are related to other proteins found throughout viral and cellular lineages.

Herpesviruses have a similar mechanism of capsid assembly as tailed bacteriophage, which has implied that Herpesviruses evolved from Caudoviruses [16]. Both classes of viruses use similar DNA packaging machinery, with a terminase-class motor and portal complex that acts as both an entrance and exit for the viral genome [17-20]. Herpesvirus and Caudovirus capsids are primarily comprised of a major capsid protein with the HK97 fold [21] and direct self-assembly using a similar scaffolding protein [22]. Herpesviruses use trimeric Triplex proteins on their coat that are positioned at the three-fold vertices of the capsid [23]. Triplex is an attractive target for HCMV vaccine development [24]. The Triplex complex is an integral part of both the immature procapsid and the final capsid shell, unlike decoration proteins of Caudoviruses [23]. Moreover, the Triplex proteins are substantially larger and share no significant sequence similarity with decoration proteins. Thus, the shared mechanisms of particle assembly and evolutionary relationships of these capsid proteins remain unclear.

Phage structural proteins can be evolutionary sources of other functions. One such example are the Anti-CRISPR proteins, which are encoded by phage and other mobile genetic elements to inhibit the CRISPR/Cas (Clustered Regularly Interspersed Short Palindromic Repeats, and CRISPR-associated system, respectively) adaptive immune system of prokaryotes [25]. Beyond their critical role in the molecular arms race between prokaryotes and viruses, Anti-CRISPR (Acr) proteins are important for medicine and biotechnology. Anti-CRISPR proteins have been found in bacterial mobile elements and have been proposed to increase virulence of these bacterial strains [26]. Acr proteins have been shown to improve CRISPR/Cas9-based genome editing by limiting off-target effects [27], as well as converting a CRISPR-associated nuclease into a transcriptional repressor [28]. Despite their importance, the evolution of Acr proteins is still largely mysterious. As has been noted previously [29,30], Acr genes are often encoded near viral structural genes, which leads us to hypothesize that some Acr genes may have evolved from structural components of viruses.

Here we investigate phage decoration protein structure, function, and evolution using a hyperthermophilic phage. Phage P74-26 is found in hot springs and infects *Thermus thermophilus* bacteria [31]. P74-26 has the longest tail of known viruses (~1µm) and thrives at 70°C [31,32], a temperature at which related mesophilic phage are disabled in minutes [33]. We identify the protein gp87 as the decoration protein of P74-26 phage. Our 1.7-Å resolution structure of gp87 reveals it has the same fold as the phage λ decoration protein, despite a surprising lack of sequence similarity. We show that the hyperthermophilic protein is substantially more stable than its mesophilic homologs. Furthermore, we identify a conserved β-barrel domain of the decoration protein that is found in Herpesvirus Triplex proteins and the Anti-CRISPR protein AcrIIC1, suggesting that these diverse proteins share a common evolutionary ancestor. Our work provides the groundwork for understanding the high stability of thermophilic viruses. Moreover, these studies illustrate a deep connection between the capsid machinery of tailed bacteriophage and Herpesviruses, and lead to a potential mechanism for the evolution of an anti-CRISPR protein.

## Results

### Major components of P74-26 virions

We purified P74-26 virions from infections of *Thermus thermophilus* strain HB8 using a combination of PEG precipitation and gradient centrifugation. The P74-26 virions consist of an icosahedral capsid ~80 nm in diameter and long, non-contractile tails exceeding 800 nm in length (Figure 1A).

**Figure 1.**
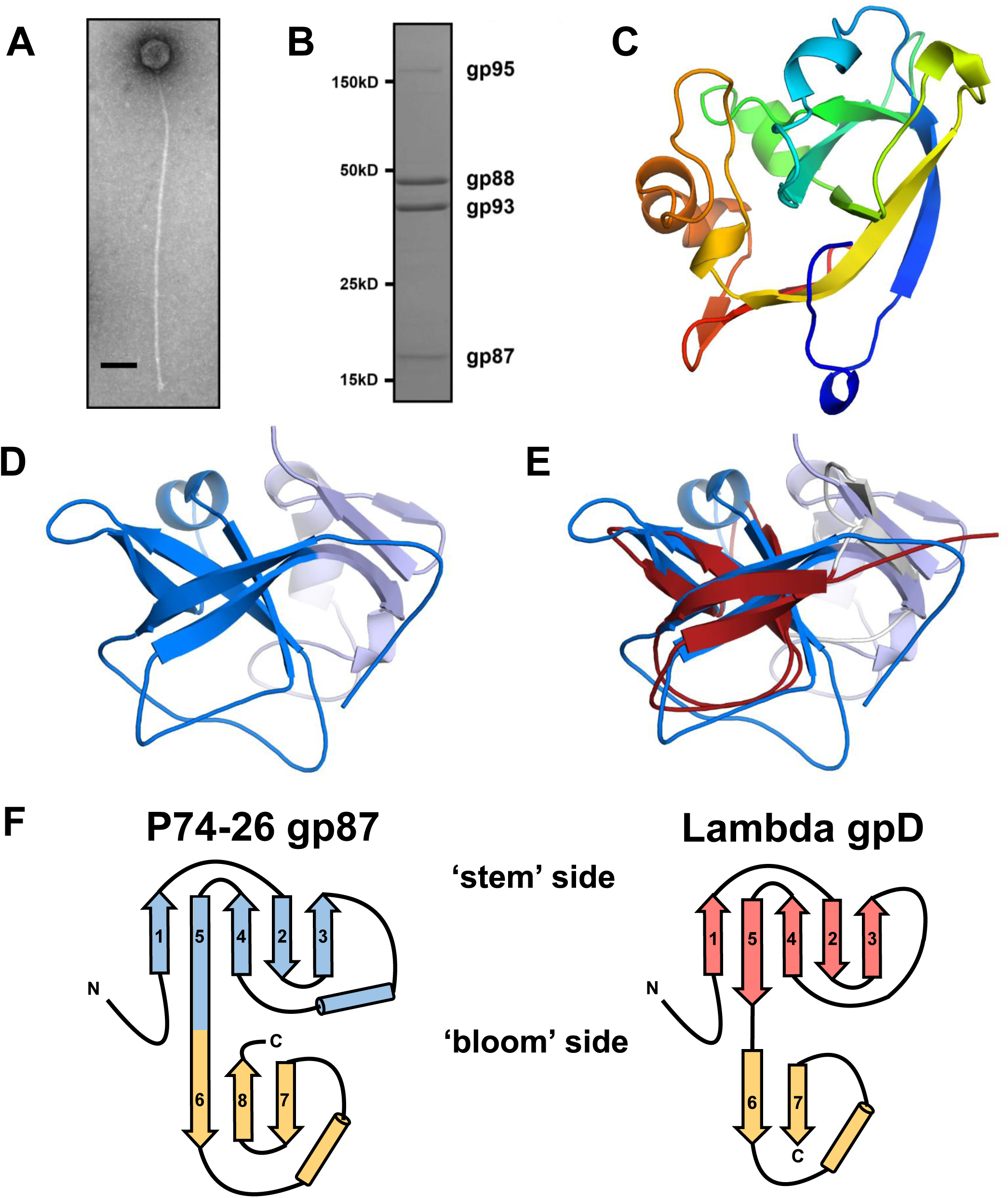
**P74-26 gp87 is a thermophilic capsid decoration protein.** A) Negative-stain electron micrograph of purified P74-26 virion; scale bar = 100 nm.
B) SDS-PAGE analysis of P74-26 virions reveals major structural components including gp87, gp88 (major capsid protein), gp93 (tail protein), and gp95 (tape measure protein).
C & D) 1.7-Å resolution structure of P74-26 gp87 (C) and with the five-stranded β-tulip domain highlighted in blue (D).
E) Structure-based alignment of P74-26 gp87 (β-tulip domain in blue) and λ decoration protein gpD (grey, β-tulip domain in red, PDB: 1C5E) reveals significant structural homology despite high sequence variance.
F) Topology diagrams of P74-26 gp87 and λ gpD reveal conserved architecture of β-tulip domain flanked by a small mixed α/β domain.

To determine whether the P74-26 capsid is stabilized by covalent interactions (as in the phage HK97 [10,34]), we analyzed the proteome of P74-26 virions by SDS-PAGE. The major capsid protein migrates on SDS-PAGE at its expected molecular weight (46.6 kDa), similar to results obtained previously (Figure 1B)[32]. This result is in contrast to phage HK97, whose MCPs are linked covalently and run on SDS-PAGE as a smear at very high molecular weight [34]. Thus, we conclude that the P74-26 capsid is stabilized by non-covalent interactions.

We hypothesized that P74-26 capsids are stabilized by decoration proteins. Our SDS-PAGE analysis reveals other major components of the virions: the tail tube protein (gp93; 37.9 kDa; the tape measure protein (gp95; 550 kDa) and gp87 (16.3 kDa), whose function in the virion is unknown. Based on size, abundance, and gene location, we hypothesize that gp87 acts as a decoration protein. gp87 is similar in size to known decoration proteins gpD and SHP from phages λ and P21, respectively (~12 kDa for both). Moreover, the relative abundance of gp87 is close to that expected for a decoration protein (318 ± 6 copies per virion by gel band densitometry). Finally, the genes encoding decoration proteins and MCPs are usually proximal to each other [35,36]; the P74-26 MCP is encoded by gene 88, adjacent to gene 87.

### P74-26 gp87 structure reveals high similarity to known phage decoration proteins

To test whether gp87 is a decoration protein, we determined its atomic structure by x-ray crystallography. Purified recombinant P74-26 gp87 readily crystallized. We solved the structure to 2.3-Å resolution by single-wavelength anomalous dispersion (SAD) using iodide ions soaked into the crystals, and phases subsequently were extended to a 1.7-Å resolution native dataset (Table 1; Figure 1C; Figure S1A). The structure reveals a core domain comprised of a five-stranded anti-parallel β-barrel, followed by a mixed α/β subdomain. According to the nomenclature for beta-strand topology [37], the barrel has a topology of (+3, +1, -2, -1) (Figure 1F). An α-helical linker connects strands 3 and 4. One end of the β-barrel is flared open, while the other is capped with loops, such that the domain resembles a tulip. We refer to the domain as a β-tulip due this resemblance. The smaller C-terminal subdomain consists of a three-stranded anti-parallel β-sheet capped by an α-helix and extended loop structure. The N-terminal 16 amino acids are not visible in the structure, most likely due to disorder. The crystal packing reveals a trimer of gp87 proteins arranged in a ‘head-to-tail’ fashion (Figure 2B). The ‘bloom’ end of the tulip is interacting with the mixed α/β-domain of a neighbor, primarily through hydrophobic interactions. β-tulip residues F27, L29, F31, I80, and L99 form an intermolecular hydrophobic cluster with the C-terminal domain residues I135 and F140 in the neighboring subunit (Figure S7C).

**Table 1.** Data collection and refinement statistics

| **Data Collection** | **KI Derivative** | **Native (APS)** |
| --- | --- | --- |
| Space Group | P 63 | P 63 |
| Wavelength | 1.54 (Home) | 0.979 (19-BM) |
| Resolution range | 29.38 – 2.34 | 25.84 – 1.69 |
| Unit cell angles | 89.74, 89.74, 31.36 | 89.52, 89.52, 31.39 |
| Unit cell dimensions | 90, 90, 120 | 90, 90, 120 |
| Total reflections | 67505 | 119377 |
| Unique reflections | 6255 | 16355 |
| Multiplicity | 10.8 (9.7) | 7.3 (7.2) |
| Completeness (%) | 98.9 (95.6) | 99.9 (100) |
| Mean I/sigma I | 16.2 (3.2) | 14.3 (2.3) |
| Wilson B factor | 32.14 | 26.76 |
| R-merge | 0.125 (0.728) | 0.057 (0.824) |
| R-meas | 0.131 (0.768) | 0.062 (0.888) |
| R-pim | 0.039 (0.242) | 0.023 (0.328) |
| CC ½ | 0.998 (0.847) | 0.999 (0.810) |
| **Refinement** |  |  |
| R-work % |  | 17.87 |
| R-free % |  | 21.26 |
| RMS bonds |  | 0.01 |
| RMS angles |  | 1.05 |
| Ramachandran favored % |  | 96.1 |
| Rotamer outliers % |  | 0 |
| Clashscore |  | 1.01 |
| Average B |  | 36.8 |

**Figure 2.**
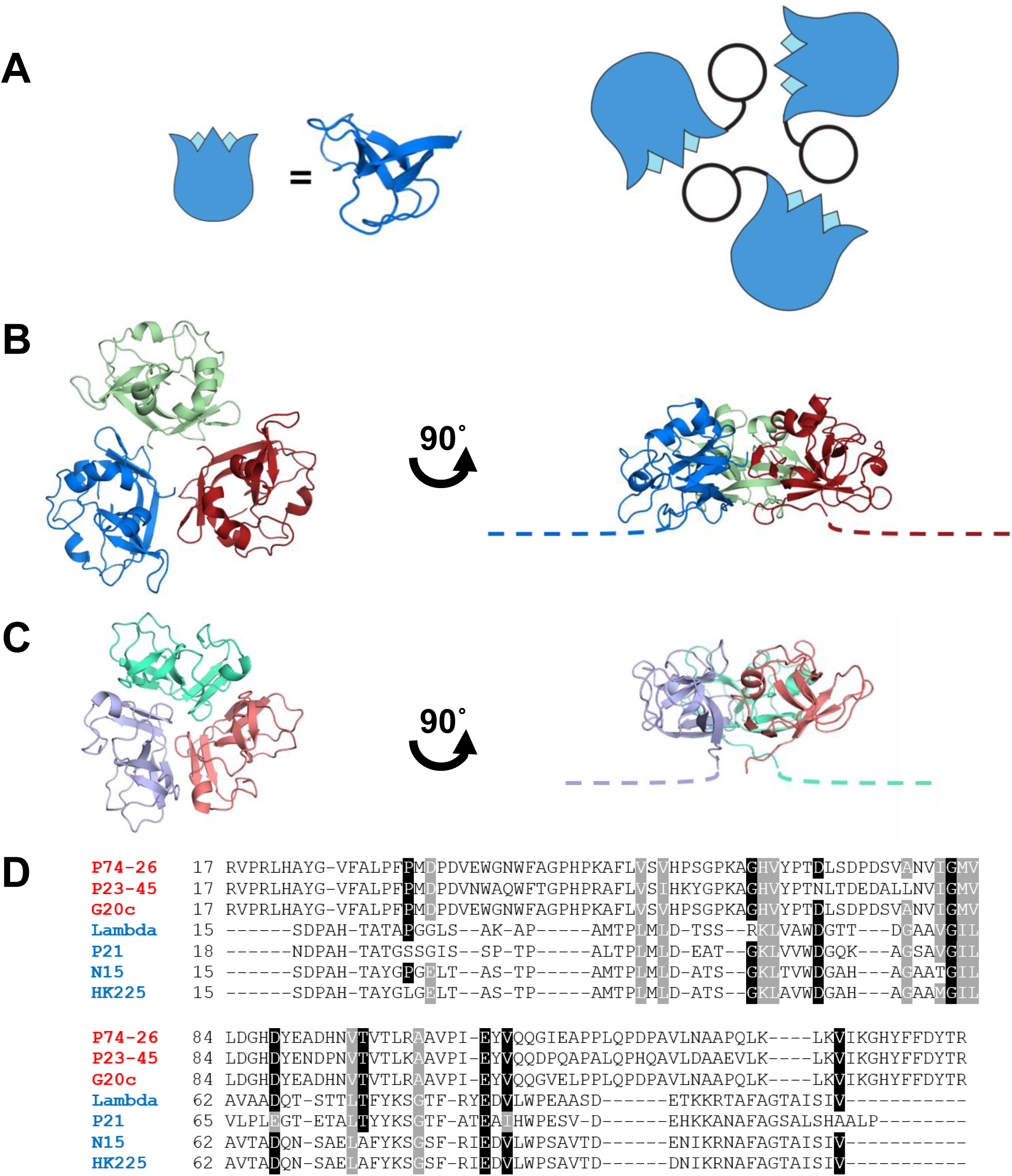
**Interactions of the decoration protein for capsid stabilization.** A. Model of decoration protein trimer highlighting positions of the β-tulip domains (blue) within the assembly.
B. P74-26 gp87 trimer highlights difference in trimer assembly characterized by a ~20° outward rotation of each of the gp87 trimer subunits (see also movie S2). The N-terminal capsid binding region of both crystal structures is disordered, and is drawn as proportional dotted lines in a and b.
C. Structure of the λ gpD trimer shows the orientation of gpD from the top of the capsid (left) and rotated 90° to the side (right).
D. Structure-based multiple sequence alignment of thermophilic and mesophilic decoration proteins reveals conserved residues. Thermophilic and mesophilic decoration proteins are color coded red and blue, respectively.

The structure and assembly of P74-26 gp87 is very similar to that observed for other well-known decoration proteins. The gp87 protein exhibits the same fold as the gpD protein from phage λ and the SHP protein from lamboid phage P21 [38,39] (Figure 1E, 1F; Figure S3A, S3B; Movie S1). Both gpD and SHP contain an N-terminal β-tulip domain and a C-terminal mixed α/β subdomain (C_α_ RMSD compared to gp87 is 2.3 Å and 2.5 Å, respectively). The similarity between the β-tulip domains of gp87 and gpD is particularly high (C_α_ RMSD = 2.1 Å; Table S1). Moreover, all three proteins have an N-terminal region that is disordered in the crystal structures (16, 14, and 11 residues for gp87, gpD, and SHP, respectively). The C-terminal domains have an overall similar fold (strand-loop-helix-strand). In gpD and SHP, the two β-strands of the C-terminal subdomain form a parallel β-sheet, whereas gp87 has an insertion of an extra β-strand that results in the formation of an anti-parallel three-stranded β-sheet (Figure S3C). Thus, the fold of gp87 is nearly identical to that of gpD and SHP.

The decoration proteins gpD and SHP also crystallized as trimers, with a similar (but not identical) arrangement around the three-fold axis of the trimer as in gp87 (Figure 2B, 2C). The trimerization interface is comprised of the core β-tulip domain of one subunit contacting the C-terminal domain of an adjacent subunit. This arrangement places the unstructured N-terminal region of each subunit at the ‘bottom’ of the trimer such that it can attach to the capsid shell [12]. Thus, the similarities in both tertiary and quaternary structure, and its high abundance in the virion lead us to conclude that gp87 is a decoration protein. The close structural and functional similarity is notable despite the low sequence homology between the thermophilic gp87 and its mesophilic cousins (~9% sequence identity) (Figure 2D).

Despite the lack of clear sequence similarity, we detect several residues that are conserved across decoration proteins. We built a structure-based sequence alignment of phage decoration proteins to identify any trends in conservation of residues throughout decoration proteins. We identify only 7 residues that are largely conserved across known homologs (Figure 2D). We mapped these conserved residues onto the λ decoration protein fold (Figure S2). The conserved residues are spread across the entire decoration protein structure.

To investigate if these ‘conserved’ residues are positioned for interaction with the capsid shell, we took advantage of the wealth of data for phage λ to model the interaction of a decoration protein with the capsid. We first positioned the gpD crystal structure into the 6.8-Å resolution structure of the phage λ capsid by cryoEM [12]. Next, we generated a homology model of the phage λ major capsid protein gpE using I-Tasser [40], which was then placed into the cryoEM density.

We note two conserved residues (Asp 66 and Glu 83 in λ gpD) which are poised near the capsid binding surface of the decoration protein and may play a role in stabilizing the capsid when gpD is bound to the virion. These residues are positioned to interact with gpE through a putative insertion in the β-hinge region and an extension of the E-loop. (We note that insertion domains are similarly present in the MCP β-hinge region of phages P22 and SPP7 [21].) Asp 66 is poised to make contact with the E-loop extension with two potential gpE binding partners (Arg 64 or Arg 66). Similarly, Glu 83 is in close proximity to the β-hinge insertion residues Lys 263 and Lys 264. Future studies will determine whether these putative interactions are important for stabilizing the capsid assembly.

### P74-26 gp87 is more stable than mesophilic homologs

We investigated whether the thermophilic decoration protein is more stable than its mesophilic counterparts. We first investigated whether gp87 forms a stable quaternary assembly in solution using size-exclusion chromatography/multi-angle light scattering (SEC-MALS). We find that gp87 is a stable trimer in solution at ~60 µM concentration, with a measured molecular mass of 53 kDa (actual molecular mass of trimer is 49 kDa) (Figure 3A). This result contrasts with λ gpD, which is monomeric in solution even up to millimolar concentrations under similar solvent conditions [38,41]. However, gpD crystallized as a trimer [38]. Thus, P74-26 gp87 forms a more stable trimer than λ gpD.

**Figure 3.**
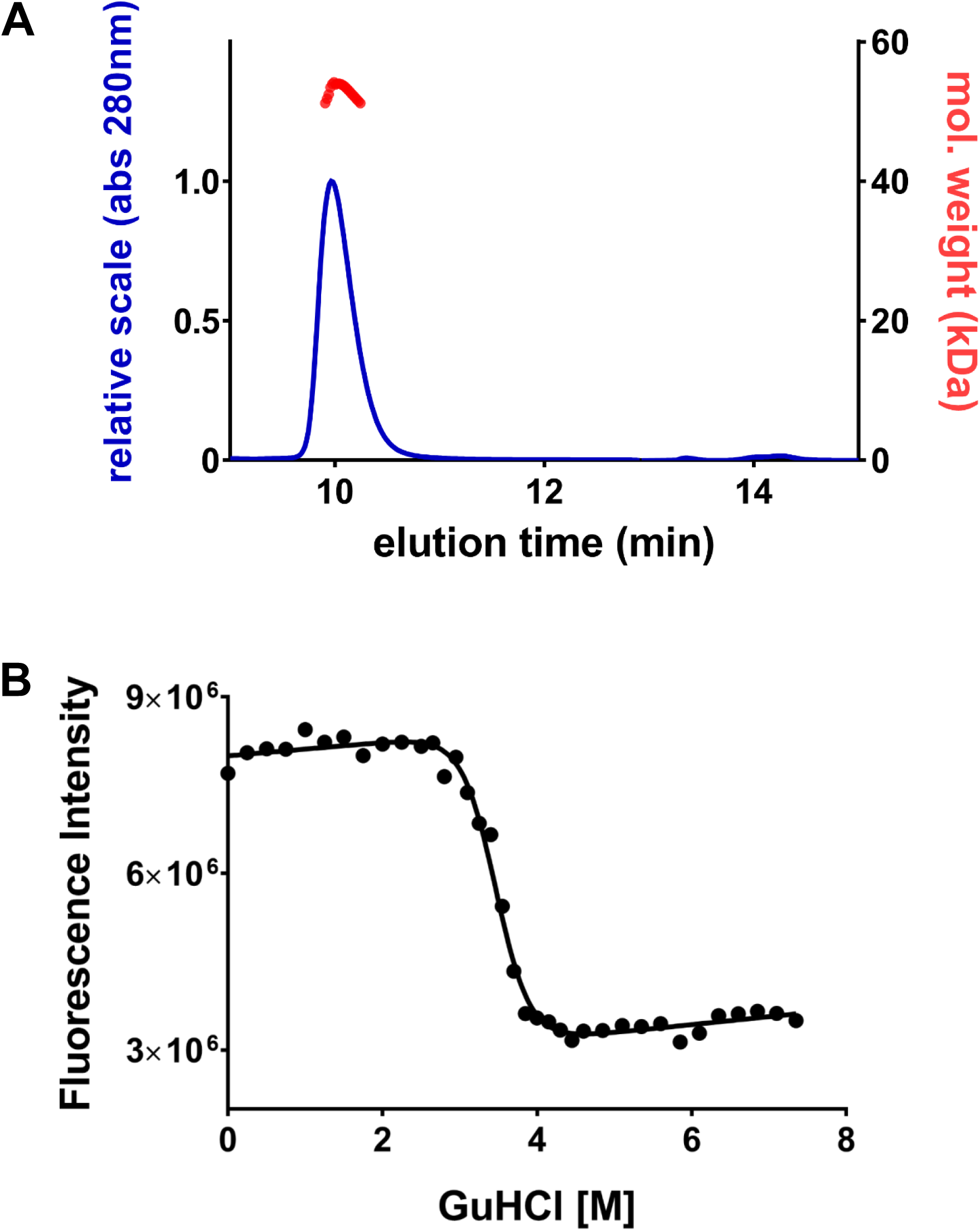
**Thermophilic decoration protein has enhanced stability compared to mesophilic homologs.** A. P74-26 gp87 forms a stable trimer in solution as determined by size exclusion chromatography-multi angle light scattering (SEC-MALS). Predicted molecular mass: 49 kDa, measured molecular mass: 52 kDa.
B. Representative equilibrium unfolding curve of P74-26 gp87 shows a steep unfolding transition from 3M to 4M GuHCl. Excitation = 295 nm, emission = 325 nm.

To investigate the comparison of protein fold stability, we measured the equilibrium unfolding energy of gp87. We induced unfolding using guanidinium hydrochloride (GuHCl). We monitored unfolding using intrinsic tryptophan fluorescence, taking advantage of the three tryptophan residues at positions 15, 39, and 42. Trp39 and Trp42 are both buried in the hydrophobic core of gp87, suggesting that these residues will report on the tertiary structure of the gp87 fold. To reach equilibrium, 0.5 μM gp87 samples were incubated at room temperature for 48 hours prior to data collection. The equilibrium unfolding curve was generated by monitoring fluorescence emission at 325 nm (Figure 3B). The unfolding curve has a steep transition between 3 M and 4 M GuHCl with a midpoint of 3.6 M GuHCl. In contrast, the midpoint of the λ gpD unfolding curve is at 1.1 M GuHCl [39], indicating that gp87 is substantially more stable than gpD. No changes in fluorescence were observed after 24 hours of incubation, indicating the samples reached equilibrium (Figure S6).

### Similarities between Herpesvirus Triplex and phage decoration proteins

Our finding that the decoration protein fold is specified with very low sequence conservation led us to investigate whether this fold is found in other proteins, but has been overlooked by classical comparative genomics approaches. We first investigated whether this ancient fold may be found in the Herpesviruses, because Herpesviruses are thought to be direct descendants of tailed bacteriophage [16,42-48]. We focused on the Triplex complex of Herpesviruses, because both Triplex and phage decoration proteins are capsid components that bind and stabilize the three-fold vertices of the capsid [12,48]. Thus, we hypothesized that the Triplex proteins and phage decoration proteins may share an evolutionary origin.

We find that core β-tulip domain of phage decoration proteins is structurally similar to the core trimerization region of the Triplex proteins (Figure 4A, 4B; Movie S3). The Triplex complex is a heterotrimer consisting of one copy of Tri1 and two copies of the Tri2 protein. Both Tri1 and Tri2 contain a cryptic β-tulip domain centrally located in their structure. In both cases, the β-tulip domains are discontinuous, with large insertions in loops between strands of the tulip (Figure 4C). The β-tulip domains of Tri1 and Tri2 have the same basic size, shape, and topology as β-tulip domains of phage decoration proteins (C_α_ RMSD = 2.4 and 2.2, respectively compared to gp87; Figure 4A-C; Table S1). The β-tulip region is the only region of obvious structural similarity between Tri1, Tri2, and decoration proteins. In all three subunits of the Triplex trimer, the β-tulip domains participate in the primary trimerization interface (Figure 4D). Moreover, the Triplex β-tulip domains are arranged roughly parallel to the capsid shell, with the bloom end of the tulip as an interaction surface, similar to the phage decoration proteins. The N-terminal regions of Triplex proteins are pointed toward the ‘floor’ of the capsid and make key interactions with the HK97 fold of MCP [48], in an analogous manner as phage decoration proteins. Both Tri1 and Tri2 contain a helical region inserted in between strands 3 and 4 of the β-tulip domain. This helical region is substantially larger in Tri2, which forms an ‘arm domain’ that tightly dimerizes the two Tri2 proteins. Insertion of helical regions is common between strands 3 and 4 of the β-tulip domain, as seen in P74-26 gp87 and other β-tulip proteins (see below). These similarities in tertiary and quaternary structure suggest a shared evolutionary history between the phage decoration proteins and the Triplex proteins of Herpesviruses.

**Figure 4.**
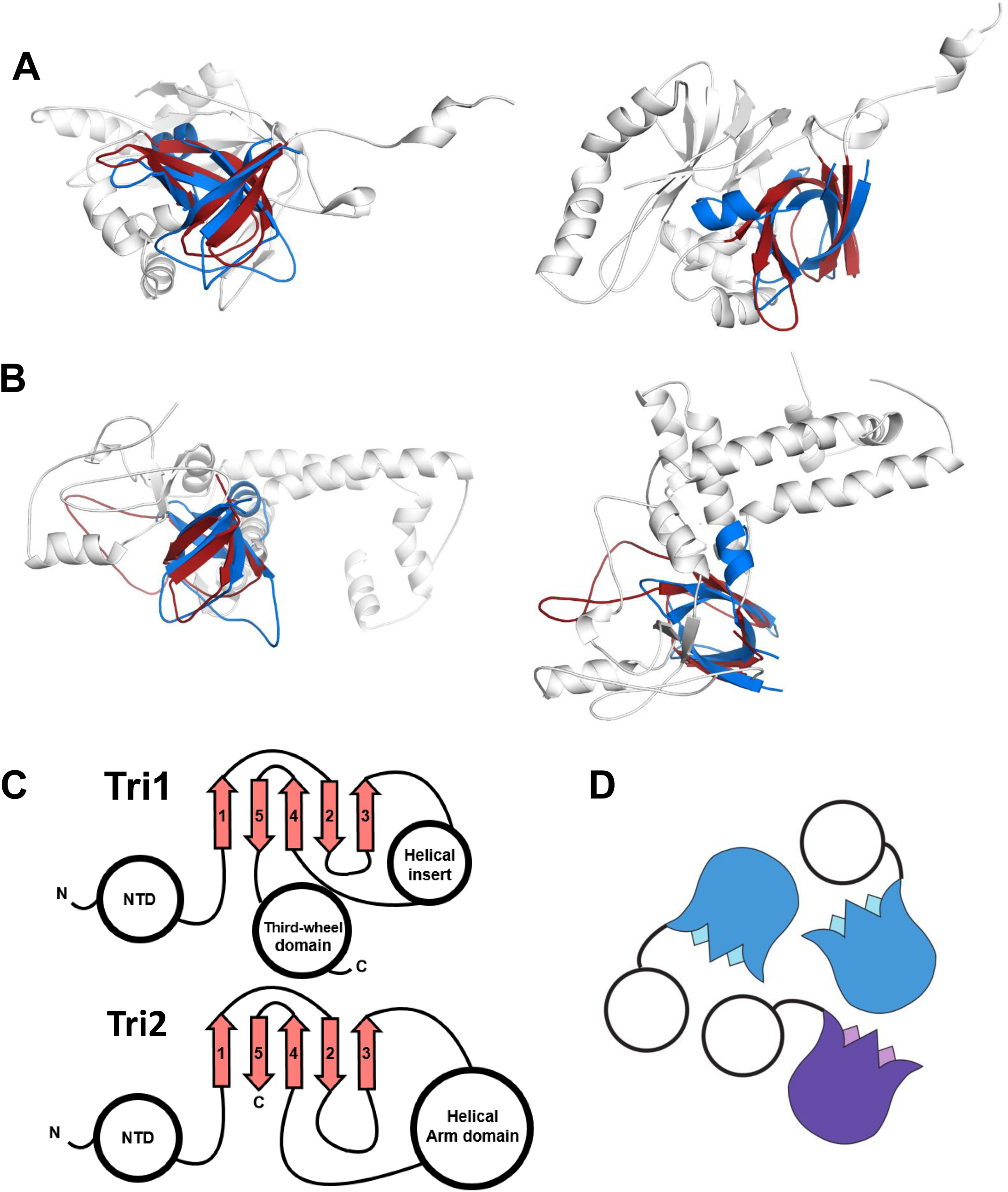
**Structural similarity of phage decoration protein trimers and the HCMV Triplex.** A & B) The P74-26 gp87 β-tulip domain (blue) is similar to central domain of HCMV Tri 1 (A) and Tri 2 (B) proteins (grey β-Tulip domains in red, PDB: 5VKU).
C) Central domain of HCMV Tri 1 and Tri2 has conserved β-tulip topology.
D) HCMV Triplex proteins form an asymmetric trimer consisting of two molecules of tri2 (blue) and one molecule of tri1 (purple).

### β-tulip domains are general protein-protein interaction motifs enriched in viruses

We sought to determine if other proteins contain β-tulip domains. Unsurprisingly, sequence similarity search algorithms such as PSI-Blast [49] found no significant homologs outside of decoration proteins or Triplex proteins. Because the β-tulip domain does not appear to be associated with a specific sequence motif, we searched for similar structures in the protein structure database using the programs DALI and VAST [50,51]. Interestingly, β-tulip domains were identified in three seemingly unrelated proteins: 1) the anti-CRISPR protein AcrIIC1 [52], 2) the tailspike protein gp12 of phage ϕ29 [53], and 3) the molybdenum metabolizing protein MoeA [54].

The anti-CRISPR protein AcrIIC1 consists of a single β-tulip with a small α-helical insertion (Figure 5A, 5B). This small protein is a broad spectrum Cas9 inhibitor and binds directly to the Cas9 HNH nuclease active site, preventing DNA cleavage [52]. Despite lacking any discernible sequence similarity, the β-tulip of AcrIIC1 is structurally very similar to that of P74-26 gp87, with a C_α_ RMSD of 2.6 Å (Figure 5A; Table S1). The β-tulip domains of both AcrIIC1 and gp87 contain a helical linker that connects strands 3 and 4 of the tulip. Unlike other β-tulip domains, AcrIIC1 is monomeric and interacts with its partner (Cas9) through the ‘stem’ end of the β-tulip (Figure 5C).

**Figure 5.**
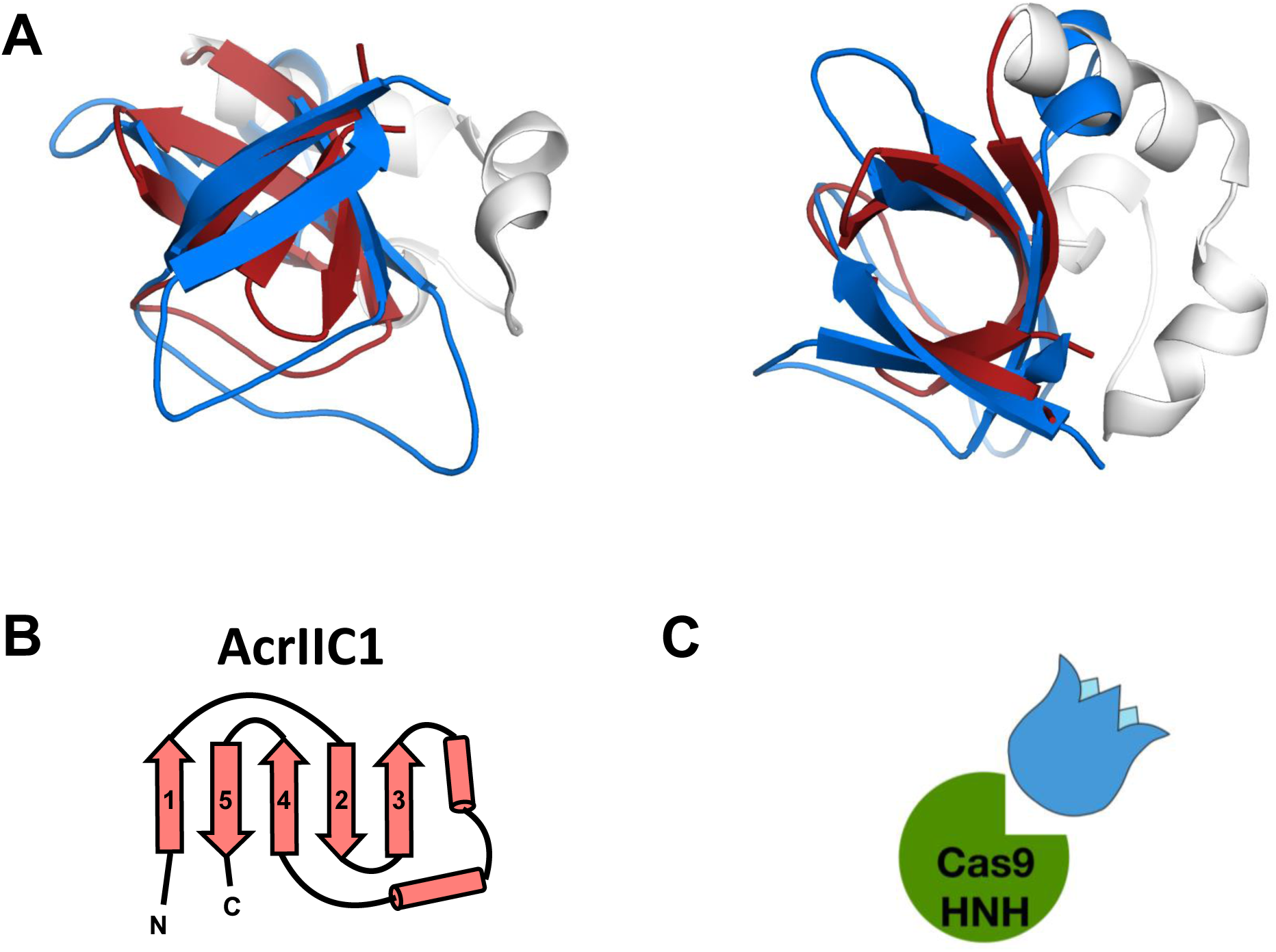
**β-tulip domain suggests evolution of anti-CRISPR proteins from phage structural proteins.** A. Structural alignment of P74-26 gp87 β-tulip domain (blue) with the anti-CRISPR protein AcrIIC1 (β-tulip in red, PDB: 5VGB).
B. Topology diagram of AcrIIC1 reveals conserved β-tulip domain architecture.
C. Cas9 binds stem side of the AcrIIC1 β-tulip domain rather than the bloom side.

The tailspike protein gp12 uses its tandem β-tulip domains (termed the D4 region) to direct assembly of the tailspike trimer [53]. The D4 region of gp12 consists of two consecutive β-tulip domains forming intramolecular contacts in a bloom-to-stem fashion (Figure S4A-D). This arrangement is echoed in the quaternary structure, as the bloom end of the C-terminal β-tulip domain interacts with the stem end of the N-terminal tulip in the adjacent subunit of the trimer (Figure S4D). The N-terminal β-tulip domain has a small β-hairpin inserted between strands 4 and 5, which projects outward away from the ring of tulips. The molybdenum biosynthetic enzyme MoeA contains a C-terminal β-tulip domain that aids in oligomerization of the MoeA trimer. As often found in other β-tulip domains, the MoeA tulip contains a short α-helical linker between strands 3 and 4 (Figure S5A, S5B).

## Discussion

### Increased stability of a thermophilic decoration protein

P74-26 is a thermophilic virus found in hot springs between 60-75° C. We sought to determine how the P74-26 virion maintains stability under these harsh conditions. We find that the P74-26 capsid is not stabilized by covalent cross-links, unlike HK97 phage that stabilizes the capsid through ‘covalent chainmail’ of the major capsid protein [10]. Instead we hypothesize that gp87 acts as a cementing protein to stabilize the capsid. gp87 is present in the virion with a stoichiometry expected for a decoration protein, and adopts similar tertiary and quaternary structure as mesophilic decoration proteins. Despite the similar structure and function, gp87 is widely divergent at the sequence level from other known decoration proteins. The low sequence homology may preclude identification of decoration proteins using classical comparative genomics approaches; structural and/or biochemical analyses may be necessary for proper annotation.

Our structural and biochemical characterization of gp87 provides insight into the mechanisms of capsid stabilization. Phage P74-26 thrives at 70° C, a temperature at which phage λ disassembles readily (t_1/2_ ~5 min)[33]. The P74-26 decoration protein is more stable than its mesophilic counterpart, gpD (Figure 3B; [39]). Furthermore, isolated gp87 is a stable trimer in solution, whereas gpD is monomeric in solution [39]. Because our unfolding data show that gp87 unfolds in a single transition, we hypothesize that dissociation of the trimer and unfolding of monomers are tightly linked. Thus, we propose that gp87 has increased stability due to both tertiary and quaternary interactions.

Our analysis of the gp87 crystal structure reveals the likely mechanism for gp87 stabilization. Each subunit in the P74-26 decoration protein is larger, burying more hydrophobic surface within each protein’s core as compared to the decoration proteins from phages λ and P21 (4470 Å^2^, 3382 Å^2^, and 3828 Å^2^ for gp87, gpD, and SHP, respectively)) [39]. In particular, gp87 displays larger clusters of branched, aliphatic residues (I, L, V, and F) than gpD (Figure S7). ILVF clusters have recently been hypothesized to be key determinants of folding and stability [55]. The larger ILVF clusters in gp87 could increase the intrinsic stability of each subunit of the trimer.

Our structure also indicates that the interfacial interactions between decoration subunits increases stability of the complex. The interfaces of the trimeric assembly bury greater surface area in gp87 than SHP or gpD. The greater buried surface area is expected to afford greater stabilization of the trimer. Again, the ILVF clusters are more extensive in thermophilic gp87 than in mesophilic gpD. gpD contains six ILVF clusters, two per subunit. In contrast, gp87 has 9 ILVF clusters, three of which span from the β-tulip domain of one subunit to the C-terminal domain of the adjacent subunit (Figure S7A, S7B). Tighter interactions between subunits will link trimerization to the unfolding of decoration protein. Indeed, our unfolding data exhibit a single transition, supporting this model.

The gp87 structure also reveals insights into how the decoration protein interacts with the capsid to stabilize the overall assembly. The N-terminal region of decoration proteins is known to interact directly with MCP, forming an extended beta-sheet across the HK97-fold E-loop [12]. This structuring of the N-terminal region creates struts that stabilize the three-fold vertices in the icosahedral shell. The N-terminal region of gp87 is substantially longer than those of mesophilic gpD and SHP (27, 19, and 21 residues long, respectively). The longer arm could provide a stronger foothold for the trimer into the capsid structure, stabilizing the three-fold vertices. Moreover, the globular domains of gp87 are larger than those of its mesophilic cousins, which could allow more interaction surface between the core trimer and the MCP shell. Furthermore, we observe that the orientation of the gp87 subunit is different than observed for gpD and SHP, with each subunit rotated downward (toward the capsid shell) by ~20° (Movie S2). The altered orientation could place the globular domains of gp87 in closer contact with the capsid shell to afford tighter contacts and greater stabilization. Finally, we identified several residues that are conserved across the known decoration proteins and we note that several of these residues appear to be poised to interact directly with the capsid shell. Interestingly, the two prime candidate residues for MCP interactions appear to interact directly with the capsid in cryoEM structures of λ phage [12]. Further structural and biochemical analyses will be necessary to uncover whether these interactions are important for stabilization of the capsid shell.

### Evolutionary relationship of Herpesvirus Triplex proteins and phage decoration proteins

Our analysis reveals that the β-tulip domain of decoration proteins is structurally and functionally conserved in Herpesvirus Triplex proteins. Although there is only 10% and 18% sequence identity between the β-tulips of P74-26 gp87 with HCMV Tri1 and Tri2, their similarities deepen our understanding of the evolutionary linkage between these two classes of viruses. The structures of the β-tulip domains are similar with the same arrangement of β-strands in the tulip. Both decoration protein and Triplex β-tulip domains participate in trimerization, in each case with the ‘bloom’ side positioned at the interface. Additionally, the β-tulip domains arrange themselves in a similar manner. Both decoration proteins and Triplex complexes position themselves at the three-fold axes of the capsid icosahedron [12,48]. In all cases, the C-terminal regions are pointed to the ‘ceiling’ of the capsid shell and the N-terminal region binds to the ‘floor’ of the capsid, with direct interactions with the E-loop of the MCP HK97 fold [12,48]. Moreover, even the variable regions appear to have significant similarities. For example, gp87, Tri1, and Tri2 have a helical region inserted between strands 3 and 4 of the β-tulip domain. In summary, the combined structural and organizational similarities illustrate the shared evolutionary history of Herpesvirus Triplex and phage decoration proteins.

For the past decade or more, evidence of the shared evolutionary history of Herpesviruses and Caudoviruses has accumulated [16]. In particular, the MCPs share the HK97 fold, although those of Herpesviruses have complicated elaborations [10,21,48]. Both virus families also direct the assembly of the capsid using a scaffolding protein that is part of procapsid but not found in the infectious particle [22,56]. Furthermore, the genome packaging machinery and mechanism is largely conserved between the two classes of viruses [17-20]. Here we add to the similarities of these different classes of viruses, showing that the decoration proteins share an evolutionary history as well. Thus, the major genes involved in virus structure and assembly are conserved throughout Caudoviruses and Herpesviruses.

While a shared evolutionary history may imply mechanistic similarities between decoration and Triplex proteins, there are substantial differences. First, the Triplex proteins are not three-fold symmetric as in phage decoration proteins (Figure 2A; Figure 4D); the two Tri2 subunits grip each other in a tight embrace, while the Tri1 protein makes less extensive contact [48]. The heterotrimeric nature of the Triplex complex has been hypothesized to allow each subunit in the complex to adopt a different role in capsid assembly [57]. Decoration proteins attach to the capsid late in the assembly process after capsid expansion [58,59], while the Triplex proteins bind the procapsid early in assembly before DNA is packaged and stay bound throughout all stages of virus particle assembly [60]. This difference in assembly may reflect the fact that the Triplex proteins embed between MCP capsomers and are an integral part of the capsid ‘floor’ [48]. This difference could be explained by the fact that the decoration proteins’ interactions with capsid components are less extensive and the decoration protein binding site is only accessible after capsid expansion [12,13].

The shared evolutionary history suggests that Triplex proteins evolved from phage decoration proteins and, over the course of time, acquired a more integral role in capsid structure. As the Triplex formed the basis for the three-fold vertices in Herpesvirus procapsids and capsids, additional domains were inserted into loops between strands of the β-tulip fold. We note that MCP also underwent domain insertions within loops of the HK97 fold during the course of evolution, suggesting a common mechanism for elaboration of the basic folds found in viral capsids. We also note that the Triplex’s role as a necessary component of the procapsid required evolution of a separate mechanism for strengthening the capsid against the internal pressure of packaged DNA [15]. In Herpes Simplex Virus 1, this is achieved by the protein UL25 [15,61], which cross-links MCP subunits within hexons [62].

### Evolutionary origin of an anti-CRISPR protein

The anti-CRISPR protein AcrIIC1 is a broad-spectrum inhibitor of Cas9 nuclease activity [30,52]. AcrIIC1 binds directly to the HNH nuclease domain of Cas9, preventing target cleavage [52]. Anti-CRISPR inhibitors of Cas9 have found use in biotechnology applications to decrease off-target DNA cleavage [27]. Despite the importance of anti-CRISPR proteins, their evolution has remained mysterious [63].

Our discovery of the structural similarity of AcrIIC1 and phage decoration proteins suggests that AcrIIC1 evolved from a phage decoration protein. AcrIIC1 consists primarily of a β-tulip fold with a strong structural similarity to the β-tulip domain of gp87 (C_α_ RMSD = 2.6 Å, Figure 5A, 5B; Table S1). However, we detect no significant sequence homology between AcrIIC1 and gp87 (6% sequence identity). This is not unexpected because the decoration proteins of λ and P74-26 phage share very little conservation, despite conserved function. Much like other β-tulips, AcrIIC1 contains a small helical insert between strands 3 and 4. Thus, the similar structures suggest that AcrIIC1 and phage decoration proteins share a common ancestral origin.

Our findings suggest a mechanism for evolution of AcrIIC1 from a decoration protein. AcrIIC1 uses the stem side of its β-tulip domain to bind the Cas9 HNH domain [52], in contrast to other β-tulip proteins that interact primarily through the ‘bloom’ side (Figure 5C). Thus, interaction with the Cas9 HNH nuclease could have evolved in a decoration protein without disrupting its central trimerization role. Gene duplication would then allow this putative bi-functional protein to specialize for each individual function (Figure 6). In support of this hypothesis, we note that Acr genes are often found near clusters of capsid genes [29]. One hurdle for this evolutionary pathway is that Acr proteins must act quickly upon phage infection to prevent CRISPR-mediated genome cleavage. However, decoration proteins are typically expressed late in infection [64] and typically remain outside the cell upon infection. How a decoration protein could be present during early stages of phage infection is not clear, although these proteins could be packaged with the genome and injected into the host cell, as is found for many phage proteins [65-68]. We surmise that AcrIIC1 could have evolved in the context of a prophage, where the Cas9 inhibitory effect could have evolved for gene regulation or other functions [28].

**Figure 6.**
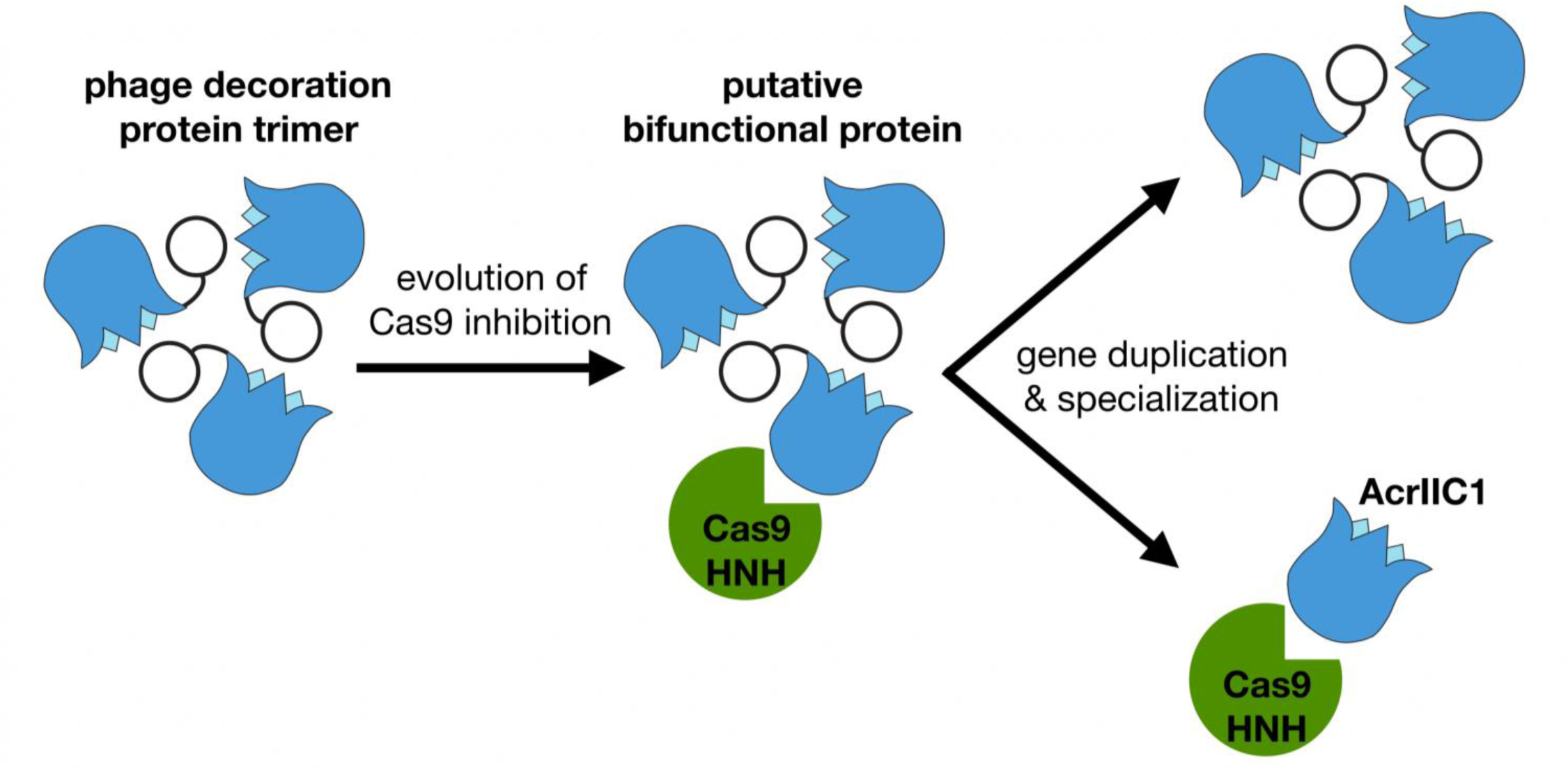
**Proposed model for the divergent evolution of decoration proteins and anti-CRISPR proteins.** Decoration protein trimers form interactions with neighboring subunits through the ‘bloom’ end of the β-tulip domain, leaving the ‘stem’ end exposed. Anti-CRISPR AcrIIC1 binds to the Cas9 HNH domain through the stem end of the β-tulip domain and may have evolved from a bifunctional phage decoration protein.

The evolution of Anti-CRISPR proteins has been mysterious. There are currently over twenty unique families of Acr proteins that have been reported [63]. These proteins are often small (50-150 residues) and have no clear sequence or structural features in common. Moreover, the mechanisms of Acr inhibition are diverse [28]. Because of their remarkable diversity and small size, Acr proteins have been proposed to evolve *de novo* from small open reading frames [63]. Here we show that the AcrIIC1 protein most likely evolved from the phage decoration protein fold. Because β-tulip domains have extremely low or no sequence homology, we could only identify this relation by structural similarity rather than sequence motifs. We propose that some Acr proteins may have evolved from other phage proteins, but their structural homology has been masked by the rapid evolution and insertion of new structural elements. Regardless, *de novo* evolution remains a likely explanation for many Acr proteins. Recent work from the Cordes group has shown that proteins from young, ‘non-coding’ genes can have rudimentary ability to fold into compact, native-like states, suggesting a mechanism for evolution of new folds [69].

### The Beta-tulip fold as a widespread protein interaction motif

Although this structural motif is quite small, we could only identify β-tulip domains in a handful of proteins. There are other examples of 5-stranded β-barrels, (e.g. HIV gp120 [70]), but these domains use a different topology than found in β-tulip domains, suggesting that they evolved independently. The β-tulip fold is structurally conserved despite lacking strong sequence conservation. This is reminiscent of the HK97 fold [21], as well as the well-studied Triose phosphate isomerase (TIM) Barrel fold [71].

The β-tulip fold is enriched in viral proteins. Of the five classes of β-tulip proteins that we identified, four of them are viral in origin: phage decoration proteins, Herpesvirus Triplex proteins, phage tailspike proteins, and phage anti-CRISPR proteins. The only outlier is MoeA, a molybdenum cofactor biosynthesis enzyme that is found throughout life. Whether the MoeA β-tulip evolved through convergent or divergent evolution is still unknown. Regardless, we propose that the viral β-tulip proteins share a common ancestor.

One of the viral β-tulip proteins is the ϕ29 phage tailspike protein, which forms a homotrimeric enzyme complex used for recognition and digestion of host cell wall structure [72]. The D4 region of the tailspike protein contains a pair of β-tulip domains arrayed in a head-to-tail fashion around the trimerization interface. Both β-tulip domains have significant structure homology to P74-26 gp87 (C_α_ RMSD = 1.7 Å and 1.5 Å for N and C-terminal β-tulip domains of D4, respectively; Figure S4A-D; Table S1). The tandem β-tulip domains in D4 act as an ‘autochaperone’ to allow trimerization of the D1, D2 and D3 regions. Removal of D4 results in non-productive assembly of the tailspike trimer [53], most likely due to kinetic traps in the folding of the highly interdigitated β-helix motifs found in the D1, D2, and D3 domains [73]. (We note that an Adenovirus uses the tailspike β-helix fold as a decoration protein to stabilize the capsid three-fold vertices [74]. Thus, both the β-tulip and β-helix regions of the tailspike have evolved to stabilize the capsid.) Interestingly, Rossmann and colleagues have noted that the D4 region of gp12 shows significant sequence homology to the minor capsid protein VP260 of *Paramecium bursaria* chlorella virus 1 (PBCV1) and some bacterial adhesion proteins [53]. The major capsid protein of PBCV1 exhibits a jelly roll fold instead of the HK97 fold found in Caudoviruses and Herpesviruses [75]. Therefore, we propose the β-tulip domain plays a role in capsid stabilization and molecular recognition across a broad swath of viral lineages, not just those of the HK97 fold viral lineage.

Our analysis reveals some general features of the β-tulip domain as a protein interaction module. The β-tulip domain tends to function in the context of a trimer (gpD, gp87, Triplex, gp12, MoeA), although AcrIIC1 is an exception. β-tulip domains act as protein-protein interaction modules, with the primary interaction surface mediated by the ‘bloom’ side of the tulip, in particular strand 1. Again, AcrIIC1 is an exception. Finally, the β-tulip motif is quite malleable in terms of sequence as well as inserted structural elements. Insertions of auxiliary elements, in particular in between strands 3 and 4 of the tulip, are common. Insertion of extra domains into the fundamental fold appears to be a common feature of capsid coat proteins; for example, HCMV major capsid protein has multiple insertions in loops of its core HK97 fold [48], and adenovirus major capsid protein has large insertions into its core jellyroll fold [75-77]. The alterations in sequence and structure of the β-tulip domain shown here provides an example of the flexibility of the fold, as well as highlights the challenges in uncovering this fold in other proteins using comparative genomics.

## Materials & methods

### Growth and purification of P74-26 virions

Phage stocks were prepared using fresh overnight cultures of *Thermus thermophilus* grown at 65° C in Thermus Growth Medium (0.8% (w/v) Tryptone, 0.4% (w/v) Yeast Extract, 0.3% (w/v) NaCl, 1 mM MgCl_2_, 0.5 M CaCl_2_). For preparation of P74-26 phage stock, 6 mL of fresh *T. thermophilus* (OD_600_ = 1.0) was inoculated with 4 mL of purified phage stock at 1x10^6^ Plaque Forming Units per mL (PFU/mL) for adsorption. Adsorption reaction mixture was incubated for 10 minutes at 65° C, then inoculated into 1 L Thermus Growth Medium. The culture was then incubated at 65° C, shaking for 4-5 hours, yielding a high-titer phage lysate (>1x10^9^ PFU/mL). Lysates were spun at 4,000 x g for 20 minutes at 4° C to remove cell debris, then the supernatant was treated with DNase I and RNase A to a final concentration of 2 Units/mL and 1 µg/mL, respectively and incubated at 30° C for one hour. Solid NaCl was added to the phage stock to a final concentration of 1 M while stirring, then culture was incubated on ice for one hour and spun at 11,000 x g for 20 minutes at 4° C. To precipitate virions, solid PEG-8,000 was added to a final concentration of 10% (w/v) while stirring and phage stock was incubated on ice overnight.

To pellet virions, precipitated phage stock was spun at 11,000 x g for 20 minutes at 4° C. The phage pellet was then resuspended in 2 mL of resuspension buffer (50 mM Tris-HCl pH 7.5, 100 mM NaCl, 1 mM MgS0_4_). 0.4 g solid CsCl was added to resuspension. Phage resuspension was then added to CsCl step gradients (five steps: 1.2 g/mL (2 mL), 1.3 g/mL (2 mL), 1.4 g/mL (2 mL), 1.5 g/mL (2 mL), and 1.7 g/mL (1 mL); made in 50 mM Tris-HCl pH 7.5, 100 mM NaCl, 1 mM MgS0_4_) prepared in 12 mL ultracentrifuge tubes (Seton Scientific). Gradients were spun in Beckman SW-40Ti swinging bucket rotor at 38,000 RPM for 18 hours at 4° C. P74-26 virions, which sediment at ~1.5 g/mL CsCl, were isolated and dialyzed twice overnight at 4° C into 2 L of 50 mM Tris-HCl pH 8.0, 10 mM NaCl, 10 mM MgCl_2_.

### SDS-PAGE analysis

30uL samples of virions (~1x10^11^ PFU/mL) were run on 12% SDS-PAGE gels. Samples were incubated in SDS loading buffer for five minutes at 95° C and run on gels at 180V for 45 minutes. Gels were fixed with 50% (v/v) ethanol, 10% (v/v) acetic acid and then stained with Coomassie Blue in 5% (v/v) ethanol, 7.5% (v/v) acetic acid. Gels were imaged on an Amersham Imager 600 (GE Healthcare). Densitometry was performed using ImageJ (Rasband, W.S., ImageJ, U. S. National Institutes of Health, Bethesda, Maryland, USA, https://imagej.nih.gov/ij/, 1997-2016).

### Electron microscopy

CsCl gradient purified virions (~1x10^11^ PFU/mL) were applied to 400-mesh copper grids (Electron Microscopy Sciences) coated in carbon. 3.5 µL samples were applied to carbon surface of grid and incubated for 30 seconds, then excess sample was removed from grids. Following sample application grids were stained with 1% Uranyl Acetate (pH 4.5) and visualized in a Philips CM120 electron microscope (120kV) fitted with Gatan Orius SC1000 camera. Micrographs were collected at a magnification of 19,500X.

### Cloning, expression, and purification of gp87

P74-26 gp87 was synthesized by the Genscript Corporation and subcloned into BamHI and XhoI sites of the pSMT3 vector with a cleavable N-terminal His6-SUMO tag [78]. Restriction enzymes were purchased from New England BioLabs, and oligonucleotide primers were obtained from Integrated DNA Technologies. P74-26 gp87 forward primer: GATCGGATCCATGGATAAAATTCAACTG; P74-26 gp87 reverse primer: GATCCTCGAGTCAGCGCGTGTAGTCAAAGAAATAG.

P74-26 gp87 was expressed in *E. coli* BLR-DE3 cells containing the pSMT3-gp87 plasmid [78]. Cultures were grown in Terrific Broth supplemented with 30 µg/mL kanamycin at 37° C to an OD_600_ of 0.7. Cultures were then incubated at 4° C for 20 minutes, then overnight expression at 18° C was induced with a final concentration of 1 mM isopropyl-β-d-thiogalactopyranoside (IPTG). Cells were then pelleted and resuspended in buffer A (50 mM Tris-HCl pH 7.5, 300 mM KCl, 20 mM Imidazole, 5 mM 2-mercaptoethanol (βME), 10% (v/v) Glycerol). All subsequent purification steps were performed at room temperature. Cells were lysed in a cell disruptor and pelleted. Cleared lysate was then applied to 2x5 mL His-Trap columns (GE Healthcare) pre-equilibrated in buffer A. P74-26 gp87 was then eluted with buffer B (50 mM Tris-HCl pH 7.5, 300 mM KCl, 500 mM Imidazole, 5 mM βME, 10% (v/v) Glycerol). Eluate was dialyzed into 50 mM Tris-HCl pH 7.5, 150 mM KCl, 2 mM DTT and the His-SUMO tag was cleaved with Ubiquitin-like Specific Protease 1 (ULP1) overnight. The dialyzed protein was passed over a 5mL His-Trap to remove cleaved His-SUMO tag. Cleaved eluate was then concentrated to 60 µM, aliquotted and flash frozen in liquid nitrogen, then stored at -80° C.

### Crystallization, data collection, and structure determination

P74-26 gp87 native crystals were formed by hanging-drop vapor diffusion at 25° C. 1mg/mL protein was mixed 1:1 with well solution containing 100 mM Tris-HCl pH 7.0, 19.5% (w/v) PEG 3350. Crystals were soaked in cryoprotectant (100 mM Tris-HCl pH 7.0, 21% (w/v) PEG 3350) and flash frozen in liquid nitrogen prior to data collection. Initial native dataset was collected to 1.9 Å using a MicroMax007-HF/Rigaku Saturn 944 CCD detector x-ray diffraction system at wavelength 1.54 Å (home source). Following native dataset collection, the native crystal was thawed, soaked in well solution supplemented with 1 M KI for 60 seconds, and flash frozen again. A KI derivative dataset was then immediately collected to 2.3 Å, with anomalous signal extending to 2.8 Å according to phenix.xtriage. Using a separate crystal, an additional native dataset was later collected to 1.7 Å at Advanced Photon Source beamline 19-BM at wavelength 0.979 Å. Diffraction datasets were processed using HKL3000 [79]. P74-26 gp87 structure was solved by SAD phasing of the 2.3 Å derivative dataset using the iodide anomalous signal with PHENIX autosol [80,81]. 11 iodides were found per monomeric subunit. Structure refinement and model building were performed with the programs PHENIX [82] and COOT [83]. Structure was deposited in the Protein Data Bank (www.rcsb.org), PDB code: 6BL5.

### Folding and refolding analysis

0.5 µM P74-26 gp87 stocks were prepared in Tris-Buffered Saline, pH 7.5 (Boston BioProducts) and either 7M Guanidinium Chloride (GuHCl) or no GuHCl. These stocks were mixed using a Hamilton Microlab 500 series titrator to yield final samples with the desired concentration of GuHCl. GuHCl concentration of stocks was determined using an ABBE Mark II refractometer (Reichert). Fluorescence emission spectra were collected at 25° C using a Fluoromax 4 Spectrofluorometer (Horiba Scientific). P74-26 gp87 was excited at 295 nm and emission spectra were collected from 310 to 400 nm for each sample. To check for hysteresis, we measured fluorescence after 24 and 48 hours of incubation at 25° C. No difference in fluorescence was seen between 24 hours and 48 hours, indicating that folding/unfolding reached equilibrium. Data was globally fit to a two-state unfolding model across all wavelengths using the in-house least squares fitting program Savuka [84]. The P74-26 gp87 unfolding was repeated in triplicate. Fluorescence intensity at 325 nm was plotted against concentration of GuHCl.

### SEC-MALS

P74-26 gp87 was run on tandem size exclusion chromatography – multi-angle light scattering (SEC-MALS) by injecting a 100 μL sample at a concentration of 1 mg/mL in 50 mM Tris-HCl pH 7.5, 150 mM KCl, 2 mM DTT. Protein was filtered through a 0.2 µM syringe filter. Elution was monitored using a Dawn Heleos-II MALS detector and Optilab T-rex differential refractive index detector (Wyatt Technology). Elution peaks were selected and molar mass was determined using the ASTRA6 analysis program (Wyatt Technology).

### Structural analysis

Homology searches using both full-length P74-26 gp87 and the β-tulip domain alone were performed using NCBI BLAST, DALI protein structure comparison server, and NCBI Vector Alignment Search Tool (VAST)[50,51]. Structure-based sequence alignment between P74-26 gp87 and λ gpD was performed using the PDBeFold program (EMBL-EBI - www.ebi.ac.uk/msd-srv/ssm/). Multiple sequence alignments with homologous proteins were performed using the Clustal Omega program (EMBL-EBI - www.ebi.ac.uk/Tools/msa/clustalo/). C_α_ RMSD and Z-score calculations for full-length protein and β-Tulip domain alignments were determined using PDBeFold. Surface area calculations for gp87, gpD, and SHP were performed using the program InterProSurf (http://curie.utmb.edu/prosurf). ILVF cluster analysis of hydrophobic networks gp87 and gpD was performed using the program BASiC Networks (http://biotools.umassmed.edu/ccss/ccssv2/basic.cgi).

## Accession Number

PDB code: 6BL5.

## Acknowledgements

The authors thank J. Hayes, C. Gaubitz, Dr. E. Sontheimer, Dr. K. Maxwell, and members of the Schiffer and Royer labs for helpful comments and discussions. The authors thank Drs. C. Gaubitz and W. Harshbarger for assistance with x-ray data collection. Drs. K. Severinov and L. Minakhin kindly provided strains of Thermus thermophilus and phage P74-26 stocks. Plasmid pSMT3 was obtained from Dr. C. Lima. The work was supported by The Pew Charitable Trusts.

## SUPPLEMENTARY FIGURE LEGENDS

**Supplementary Figure 1. P74-26 gp87 structure solved using iodide phasing.**

Iodide ion binding positions are shown mapped onto 1.7-Å structure of gp87 (iodides in grey).

**Supplementary Figure 2. Conserved residues in phage decoration proteins make contact with the capsid surface.**

Conserved residues mapped onto λ decoration protein gpD (blue) and fit into the EM density of the λ virion (EMBD: 5012). A homology model of λ major capsid protein (gpE) was likewise fit into the capsid EM density (purple and green). Highlighted residues Asp 66 and Glu 83 are positioned toward the capsid surface and may make critical contacts with the capsid shell.

**Supplementary Figure 3. P74-26 gp87 has significant similarity to known decoration proteins.**

A. Structural alignment of gp87 (β-tulip in blue) with the decoration protein SHP from phage P21 (white, β-tulip in red, PDB: 1TD3).
B. Topology diagram of the P21 decoration protein SHP.
C. Structural alignment of P74-26 gp87 (blue) with λ gpD (grey) highlighting additional strand in gp87 (red) forming an antiparallel β-sheet in the C-terminal domain.

**Supplementary Figure 4. Structural similarity with phage tail spike proteins.**

A. Structure-based alignment of P74-26 gp87 β-tulip domain (blue) with the N-terminal β-tulip domain (red) of the D4 domain of Phi29 tail spike protein gp12.
B. Alignment of P74-26 gp87 β-tulip domain with the C-terminal β-tulip domain of gp12.
C. Topology diagram of Phi29 gp12 shows the orientation of the tandem β-tulip domains.
D. Model of the head-to-tail β-tulip domain orientation in gp12.

**Supplementary Figure 5. Conservation of β-tulip domain in the Moco biosynthesis enzyme MoeA.**

A. Structure-based alignment of P74-26 β-tulip domain (blue) with MoeA (grey, β-tulip in red, PDB: 2NQQ).
B. Topology diagram of MoeA E-domain.

**Supplementary Figure 6. gp87 GuHCl titrations reach equilibrium after 24 hours.**

Equilibrium unfolding of gp87 after incubation for 24 hours (black), 48 hours (red), and 120 hours (blue) (independent experiments) shows equilibrium is reached after 24 hours. Excitation = 295 nm, emission = 325 nm.

**Supplementary Figure 7. P74-26 gp87 trimer forms an extensive hydrophobic network.**

A. Hydrophobic ILVF clusters mapped onto the structure of λ gpD trimer show clusters within the β-tulip and C-terminal domains that are unconnected to each other.
B. ILVF clusters mapped onto the P74-26 gp87 trimer show an extended hydrophobic network, with clusters forming intermolecular and intramolecular interactions (clusters shown in blue span multiple subunits).
C. gp87 β-tulip domains interact with neighboring subunits primarily through hydrophobic interactions, forming an intermolecular cluster consisting of Ile (red), Leu (green), and Phe (blue) residues.

**Supplementary movie 1.**

Structure-based alignment of gp87 (blue) and gpD (red) shows significant structure conservation despite having low sequence homology.

**Supplementary movie 2.** Changes in subunit orientation between gp87 and gpD. gp87 trimer orientation is characterized by a 20° outward rotation of individual subunits relative to the position of the subunits in the λ gpD trimer.

**Supplementary movie 3.** Similarities of conserved β-tulip domains in gp87, Tri1, Tri2, and Anti-CRISPR AcrIIC1.

**Table S1.** C_α_ RMSD comparison of β-tulip domains

